# Delivery of cell-penetrating chromatin sensor-actuators to human osteosarcoma cells

**DOI:** 10.1101/2020.02.28.969907

**Authors:** Stefan J. Tekel, Nicholas Brookhouser, Karmella A. Haynes

**Affiliations:** School of Biological and Health Systems Engineering, Arizona State University, Tempe, AZ 85287, USA; Clinical and Translational Sciences Graduate Program, University of Arizona College of Medicine-Phoenix, Phoenix, AZ 85004, USA; Wallace H. Coulter Department of Biomedical Engineering, Emory University, Atlanta, GA 30322, USA

## Abstract

The leading FDA-approved drugs for epigenetic cancer therapy are small molecule compounds that activate silenced tumor suppressors by inhibiting enzymes that generate aberrant repressive chromatin. Although promising, this approach is limited because chromatin-modifying enzymes often target non-chromatin proteins and can serve dual functions as gene repressors and activators. Previously, we have demonstrated that a transgenically expressed synthetic polycomb-derived transcription factor (PcTF) could activate genes in silenced chromatin via specific interactions with histone H3 trimethylated at lysine 27 (H3K27me3). Efficient non-viral intracellular delivery remains a challenge for protein-based biologics. Herein, we report the delivery of cell-penetrating PcTF (CP-PcTF) to cultured cells. We expressed and purified recombinant PcTF that was fused in frame with a nuclear localization signal and a cell penetrating peptide tag (TAT). We demonstrated rapid and efficient uptake of soluble CP-PcTF by osteosarcoma U-2 OS cells grown in 2-D monolayers and 3-D spheroids. However, CP-PcTF had a modest effect on gene expression and cell proliferation compared to transgenically-expressed PcTF from our previous work. Overall, these results suggest that TAT is a very effective delivery vehicle for the recombinant transcriptional regulator PcTF, and that further technical development is needed to deliver functional PcTF into cell nuclei.

## INTRODUCTION

Molecular pathways that support healthy and diseased cell states are in large part driven by proteins that act outside and within cells. Therefore, protein-based therapeutics (biologics) offer potential advantages over small molecule drugs and gene therapy, such as high specificity, potency, and lower likelihood to induce immune responses [1,2]. Recombinant protein delivery avoids nucleic acid induced cytotoxicity and off target effects from viral or DNA particles. In addition, nucleic acid based delivery can be cytotoxic and induce recombination with the endogenous genome. Recombinant protein delivery avoids these caveats and has been shown to enhance the specificity of targeted interventions. For instance, a Cas9 base editor (HF-BE3) delivered to HEK293 cells as ribonucleoproteins specifically edited target cytosines, whereas delivery of HF-BE3-expressing plasmid DNA led to off target editing [3].

Clinical biologics such as antibodies [4] and insulin [5] interact with the exterior of the cell or act within the cytoplasm. However, many critical targets, such as genes and epigenetic factors, lie within the nucleus. Pre-clinical, basic research has demonstrated nuclear delivery of Cas9/gRNA gene-editing complexes in cultured animal cells via cationic lipids [6] or nucleofection [7], in tissues *in vivo* via cationic lipids [6] or local injection [7], and in plant cells via PEG-calcium [8]. Additionally, fusions of protein cargos with cell penetrating peptides (CPPs) have been successfully used to deliver metalloenzymes to catalyze the release of small molecules in cells [9], transcription factor HOXB4 to expand stem cells [10], and p53 to inhibit cancer cell growth [11]. The CPP induces endocytosis when the fusion molecule reaches the cell membrane [12]. CPPs such as TAT from HIV are small (11 amino acids), are unlikely to interfere with protein function [13,14], and therefore could be used for a variety of biologics, including genome- and epigenome-targeting proteins.

We are interested in developing cell-penetrating protein biologics for epigenetic therapy. Currently, the leading epigenetic drugs are small molecule compounds that block enzymes that modify chromatin, and can result in undesired effects, such as oncogene activation [15]. Engineered chromatin proteins could provide greater target specificity. For instance, epigenome editing proteins have been used to alter disease-associated histone post-translational marks and DNA methylation at specific genes and enhancers (reviewed in [16,17]). We have previously designed chromatin sensor-actuator proteins that bind histone post-translational modifications and co-activate groups of repressed genes. We demonstrated that a H3K27me3-specific sensor-actuator called PcTF activates epigenetically repressed tumor suppressor genes in several cancer cell lines [18,19]. However, experiments with epigenome editors and sensor-actuators have relied heavily on the delivery and overexpression of fusion-encoding transgenes. Having recently demonstrated that purified recombinant PcTF specifically binds its target H3K27me3 *in vitro* [20], we set out to determine if purified PcTF proteins maintain their function as potent gene activators after they are delivered directly to cells.

To determine which peptides would enable delivery of PcTF, we tested CPPs with different cellular trafficking properties, and analyzed mCherry-tagged PcTF uptake in cultured osteosarcoma U-2 OS with fluorescence microscopy and flow cytometry. In this report, we describe the results of co-delivery of purified PcTF with one of four CPPs: HA2 [21], E5 [22], TAT [1], L17E [23]. TAT and L17E peptides showed the highest levels of cellular mCherry signal after 24 hours of treatment. In order to streamline protein delivery and reduce excess unbound CPP’s, we fused a C-terminal TAT peptide in-frame with PcTF to produce CP-PcTF. We observed that CP-PcTF accumulates in cells in a dose and time-dependent manner. Changes in gene expression in CP-PcTF-treated cells suggest that the protein remains active after it enters the cytoplasm, and eventually the nucleus. In addition, we show that CP-PcTF slows the growth of U-2 OS spheroids, suggesting that Cp-PcTF-mediated gene regulation is sufficient to affect the phenotype of these cells. These results represent important progress towards the nuclear delivery of recombinant chromatin proteins to cells in 2D and 3D models.

## RESULTS

### Identification of cell-penetrating peptides that support PcTF delivery

In order to identify an effective protein delivery vehicle for PcTF, we tested four different CPPs (Figure 1; Supplemental Table S1, Figure S1). TAT is a peptide fragment from the human immunodeficiency virus 1 (HIV-1) that induces pinocytosis and shows robust cell penetrating ability [21]. Cargo delivered by pinocytosis can become trapped in endosomes, leading to loss of protein function [24] and failure to enter the nucleus. Therefore, we also tested two TAT hybrid peptides that carry signals to support endosomal escape. TAT-HA2 includes the N terminal 20 amino acids of the influenza virus hemagglutinin protein [21]. E5-TAT includes an enhanced HA2 with increased membrane disruption activity [22]. We also tested an engineered membrane-lytic peptide called L17E, which was previously shown to enable delivery of protein cargo [23]. We hypothesized that PcTF would enter the cell through macropinocytosis along with the CPP’s, which contain positively charged residues that interact with the negatively charged cell membrane to induce cell entry. We presumed that once in the cell, PcTF would escape the endosome and be imported into the nucleus where it can bind to H3K27me3 sites and upregulate epigenetically repressed genes (Figure 1).

**Figure 1.**
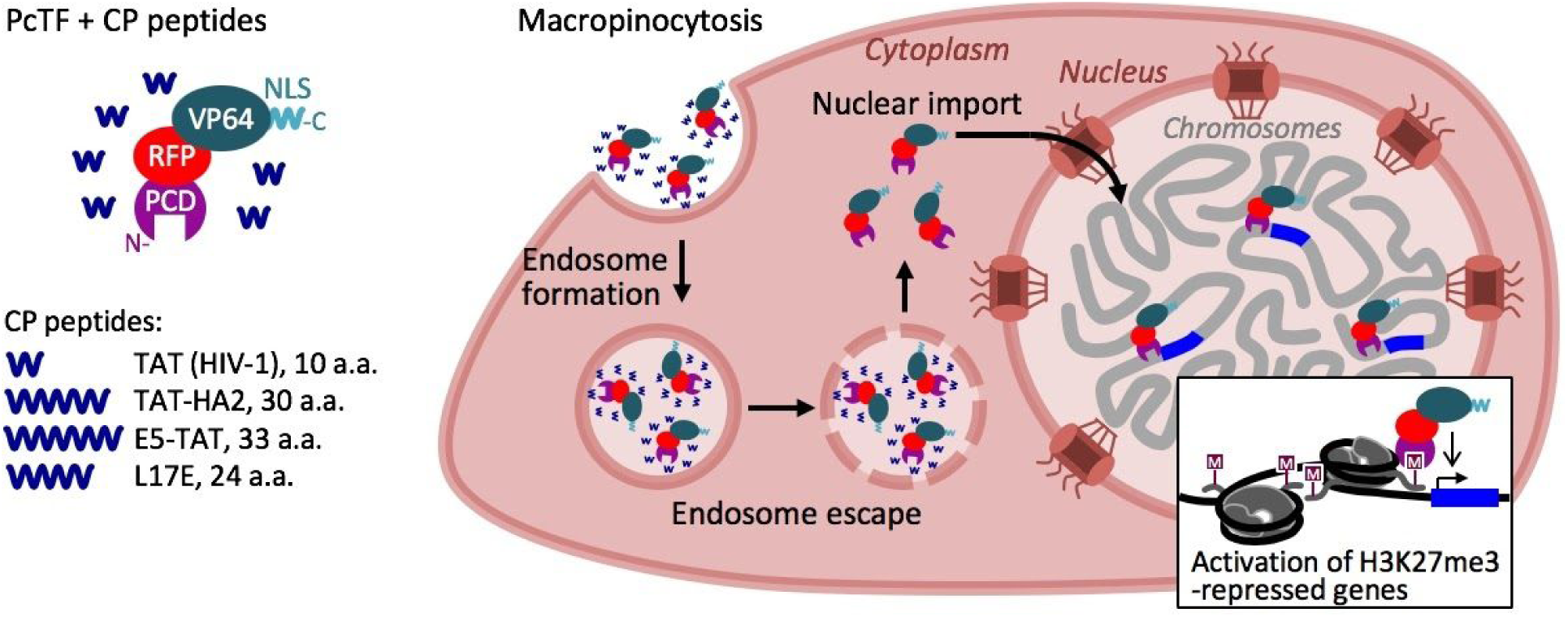
Schematic for the delivery and cellular transport of the fusion chromatin protein PcTF. Purified PcTF is incubated with a small cell-penetrating peptide (TAT, TAT-HA2, E5-TAT, or L17E) and added directly to the growth media in a well of cultured U-2 OS cells. PcTF enters the cell through macropinocytosis, followed by endosomal escape into the cytoplasm. PcTF is then imported into the nucleus where it activates H3K27me3-silenced genes.

We performed co-delivery of purified PcTF and each cell-penetrating peptide. First, we incubated each peptide (5.0 μM) with a high concentration (1.0 μM) of recombinant PcTF for 24 hours at 37°C. We added these preparations to U-2 OS cells that were grown to 60% confluency and allowed protein uptake to occur for 24 hours. Red fluorescent protein (RFP) signal from the mCherry domain within the PcTF fusion protein allowed us to visualize cellular uptake. Cells were washed three times with PBS before imaging with fluorescent microscopy. PcTF without cell-penetrating peptide showed very little detectable RFP compared to untreated cells. In contrast, PcTF that was incubated with TAT or L17E showed strong RFP signal within the boundaries of the cell cytoplasm (Supplemental Figure S1). Most of the RFP signal from the PcTF/TAT-HA2 and PcTF/E5-TAT-treated samples appeared as extracellular aggregates that did not coincide with individual cells. These PcTF complexes may have aggregated before entering the cells, or induced lysis of PcTF-containing cells. Next, we set out to determine if cell-penetrating activity could be integrated into the PcTF protein by fusing TAT or L17E peptides to the C-terminus of PcTF.

### The TAT peptide signal supports delivery of purified PcTF into U-2 OS cells in 2-D and 3-D cultures

We constructed two fusions for recombinant expression and purification: PcTF-TAT and PcTF-L17E. We hypothesized that these fusion proteins, generally referred to here as cell-penetrating PcTF (CP-PcTF), would enter the cell through macropinocytosis in a similar manner as smaller cell-penetrating peptides (Figure 1). To obtain sufficient quantities of purified protein for cell delivery, we added a 6-histidine tag to the N-terminus for nickel column purification. We cloned each CP-PcTF open reading frame in frame with the 6-histidine sequence in an isopropyl β-D-1-thiogalactopyranoside (IPTG) inducible pET28 vector and transformed each recombinant plasmid into *E. coli* for IPTG-induced overexpression. Only the TAT fused PcTF was fully expressed in *E. coli*, as indicated the presence of red color from mCherry, and a prominent 46 kD band visualized by PAGE of whole-cell lysates (data not shown). Therefore, we continued our study using only PcTF-TAT. Lysates from pelleted PcTF-TAT-expressing cells were loaded into a Nickel-NTA binding column. The eluted proteins were concentrated into phosphate buffered saline (PBS) and analyzed for purity by polyacrylamide gel electrophoresis (PAGE) (Supplemental Figure S2) prior to cell treatments. In addition to the expected product (∼46.2 kDa), we observed two additional smaller bands (Supplemental Figure S2) that suggested cleavage of a portion of the protein, which has been observed for overexpressed recombinant proteins in certain *E. coli* strains [25]. Cultured U-2 OS cells treated with 0.2 μM purified CP-PcTF showed much stronger cellular red fluorescent signal after 24 hours than cells treated with PcTF that lacked the TAT signal (Figure 2A). This result indicated that TAT enabled more efficient protein uptake as a C-terminal signal fused to PcTF.

**Figure 2.**
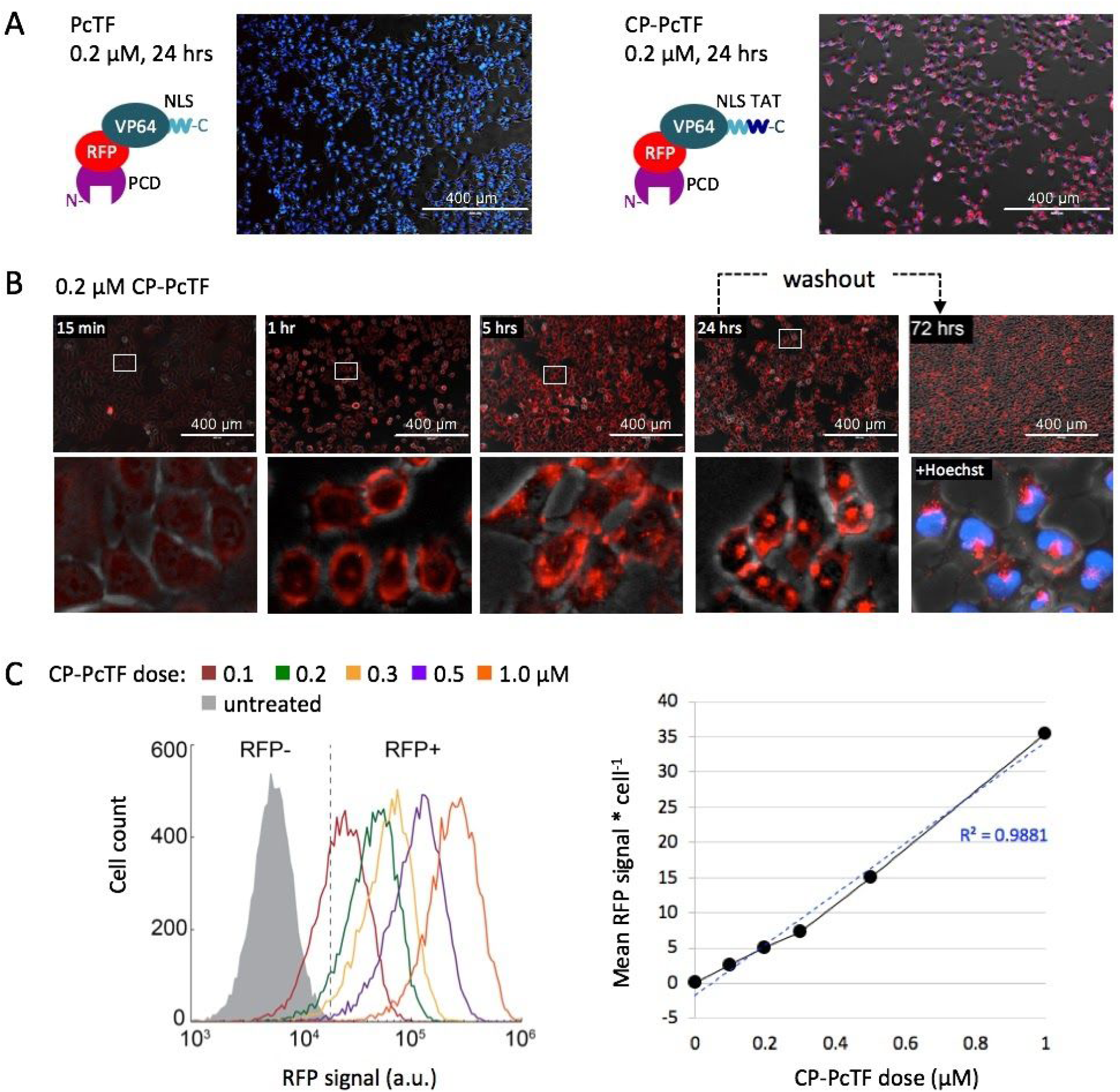
Uptake of soluble CP-PcTF by U-2 OS cells grown as two-dimensional (2-D) a monolayer. (A) Comparison of uptake of 200 nM soluble PcTF versus CP-PcTF. Each fluorescence microscopy image shows merged mCherry (red), Hoechst nuclear DNA stain (blue) and phase contrast channels. (B) Fluorescence microscopy was used to observe cellular uptake of CP-PcTF over time. Cells were treated with 200 nM of CP-PcTF for 15 minutes, 1 hour, 5 hours, or 24 hours, washed three times with PBS, and imaged. The 24 hour sample was cultured for an additional two days after washout (dotted arrow), stained with Hoechst nuclear dye, and imaged again. (C) Flow cytometry was used to quantify red fluorescent protein (RFP) signal from U-2 OS cells treated with 0.1 - 1.0 μM CP-PcTF for 24 hours. The histogram shows RFP signals from ∼10,000 live cells (per sample) that were gated by forward and side scatter. The x-y dot plot shows the mean RFP signal divided by the live cell count for each population.

Next, we carried out a time-course treatment to determine the earliest point at which CP-PcTF became completely internalized by the cells. U-2 OS cells were grown as a monolayer to 75% confluency, treated with 0.2 μM soluble, purified CP-PcTF proteins for 15 minutes, 1 hour, 5 hours, or 24 hours. At the end of each time point, extracellular protein was washed away with phosphate buffered saline (PBS), and cells were imaged via fluorescence microscopy (Figure 2B). We observed the strongest mCherry signal at the edges of each cell (presumably the cell membrane) in samples that had been treated for 15 minutes or 1 hour. In the sample that was treated for 24 hours the strongest RFP signal appeared at the center of each cell. We continued to culture the sample that was treated for 24 hours for an additional two days. Similar levels of RFP signal persisted within the cells at the 72 hour time point. This result suggests that CP-PcTF is taken up by cells through regulated endocytosis instead of diffusion (Figure 2B). To determine the nuclear uptake of CP-PcTF, we repeated the 24 hour treatment, followed by two days of growth after washout, and stained the cells with Hoechst (a DNA-specific dye) to image the position of RFP signal relative to the nuclei. RFP signals appeared as discrete particles or larger clusters in or near each nucleus.

To further investigate the kinetics of CP-PcTF uptake, we treated cells with increasing concentrations of CP-PcTF. Since TAT-stimulated macropinocytosis is a continual non-selective process, we expected that the amount of internalized CP-PcTF would scale with extracellular concentration (dose). We used a concentration range based on other reports that demonstrated successful delivery of 2.0 nM to 5.0 μM TAT-fused proteins to cultured cells [10,13,14,26]. We treated U-2 OS cells with 100 nM to 1.0 μM CP-PcTF for 24 hours, washed out excess extracellular protein, and harvested the cells for flow cytometry. We observed a unimodal distribution of live, RFP-positive cells for all treatments, with a clear shift in signal intensity at least one order of magnitude higher than the untreated control sample (Figure 2A). The mean RFP signal intensity increased linearly with CP-PcTF dose concentration (R^2^ = 0.99). These results demonstrate that the amount of intracellular CP-PcTF can be regulated by the concentration of CP-PcTF that is added to the media during a 24-hour treatment.

### U-2 OS spheroids show delayed growth after treatment with CP-PcTF

The spatial organization of cells in three dimensional (3-D) tumor models or “spheroids” more closely reflect cancerous tissues than 2-D cultures [27], and are therefore important for understanding the activity of potential therapeutics. We set out to investigate the uptake of CP-PcTF in U-2 OS spheroids, and to determine the impact of CP-PcTF treatment on cell proliferation. We generated spheroids by culturing cells in low attachment plates with constant, gentle rotation for three days. We treated the spheroids with CP-PcTF for 24 hours and used fluorescent microscopy to visualize cellular uptake of the protein (Figure 3). We observed a general overlap of RFP signal from CP-PcTF with DNA stain (Hoechst) in spheroids (Figure 3A), as we had observed in the 2-D cultured cells (Figure 2B). To quantify CP-PcTF uptake, we dissociated the spheroids and measured RFP in single cells via flow cytometry. In contrast to the 2-D cell cultures, cells from spheroids showed a broad range of no signal to high levels of RFP signal, indicating greater variability of CP-PcTF uptake by individual cells (Figure 3B). For the highest concentration of CP-PcTF (1.0 µM), we observed 45.4% RFP-positive cells from spheroids, versus 100% from 2-D cultures. Our results suggest that CP-PcTF does not completely penetrate all cells within spheroids after 24 hours of treatment. Uptake appeared to scale linearly (R^2^ = 0.92) with CP-PcTF concentration, indicating that delivery might be improved with higher doses.

**Figure 3.**
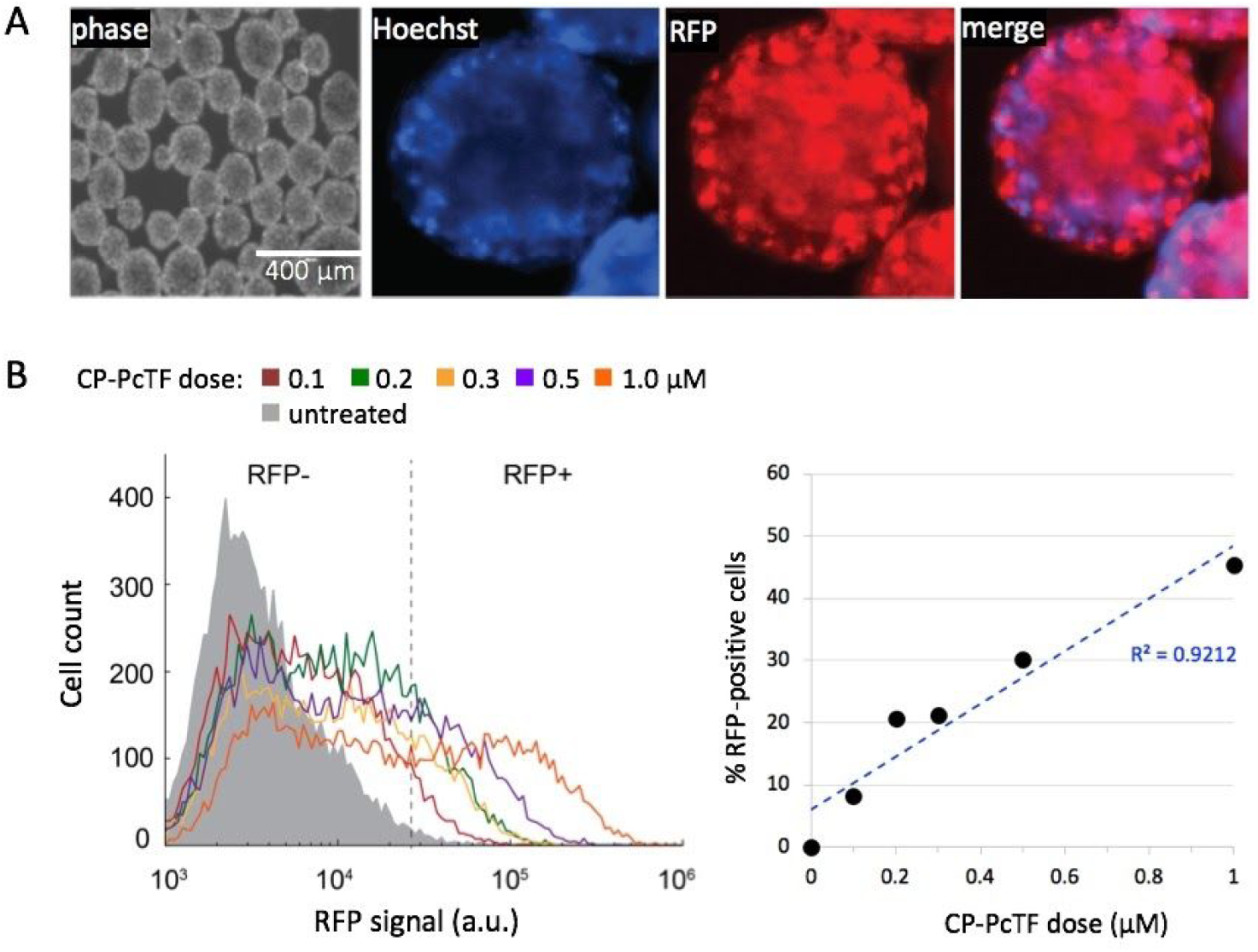
Uptake of CP-PcTF in U-2 OS spheroids. (A) U-2 OS spheroids were grown up to 40 µm in diameter before treatment with CP-PcTF for 24 hours. Wide field fluorescence microscopy showed general overlap of RFP signal from CP-PcTF with DNA stain (Hoescht). A representative spheroid is shown here. (B) Spheroids were treated with 0.1 to 1.0 µM CP-PcTF for 24 hours, collected and dissociated with accutase, and resuspended in 1x PBS for flow cytometry analysis. The histogram shows RFP signals from ∼10,000 live cells (per sample) that were gated by forward and side scatter. The x-y dot plot shows the the percentage of RFP-positive cells for each sample.

To investigate how CP-PcTF might affect the proliferation of cells within tumor tissue, we measured spheroid growth following treatment with CP-PcTF. In a previous study, we observed that over-expression of PcTF in U-2 OS cells increased senescence-associated beta-galactosidase levels and reduced cell proliferation (colony size) in 2-D U-2 OS cultures [28]. Therefore, we hypothesized that CP-PcTF might inhibit the growth of U-2 OS spheroids. We treated spheroids with 1.0 μM CP-PcTF for 24 hours, washed away excess protein, incubated the spheroids in fresh growth medium, and measured the diameters of 100 spheroids per sample for up to 10 days. These and subsequent experiments included CP-mut, which carries a deletion of the N-terminal H3K27me3-targeting domain (PCD), as a negative control.

On day four, we observed a significant reduction in the spheroid diameters for the CP-PcTF treated sample compared to the untreated sample (*p* = 8.3E-10) and the sample treated with CP-mut (negative control protein) (*p* = 1.6E-13) (Figure 4A). RFP signal remained visible throughout the spheroids (Figure 4B), indicating that at least some portion of the cells retained PcTF after cell division. On days five, six, and ten, the mean diameter of CP-PcTF-treated spheroids continued to increase relative to day four, but remained significantly lower than the two negative control samples (*p* < 0.001). Therefore, a transient 24-hour treatment with CP-PcTF (followed by washout) might temporarily hinder overall spheroid growth, while some of the cells within each spheroid might continue to proliferate. We consider possible underlying causes of this outcome in the Discussion.

**Figure 4.**
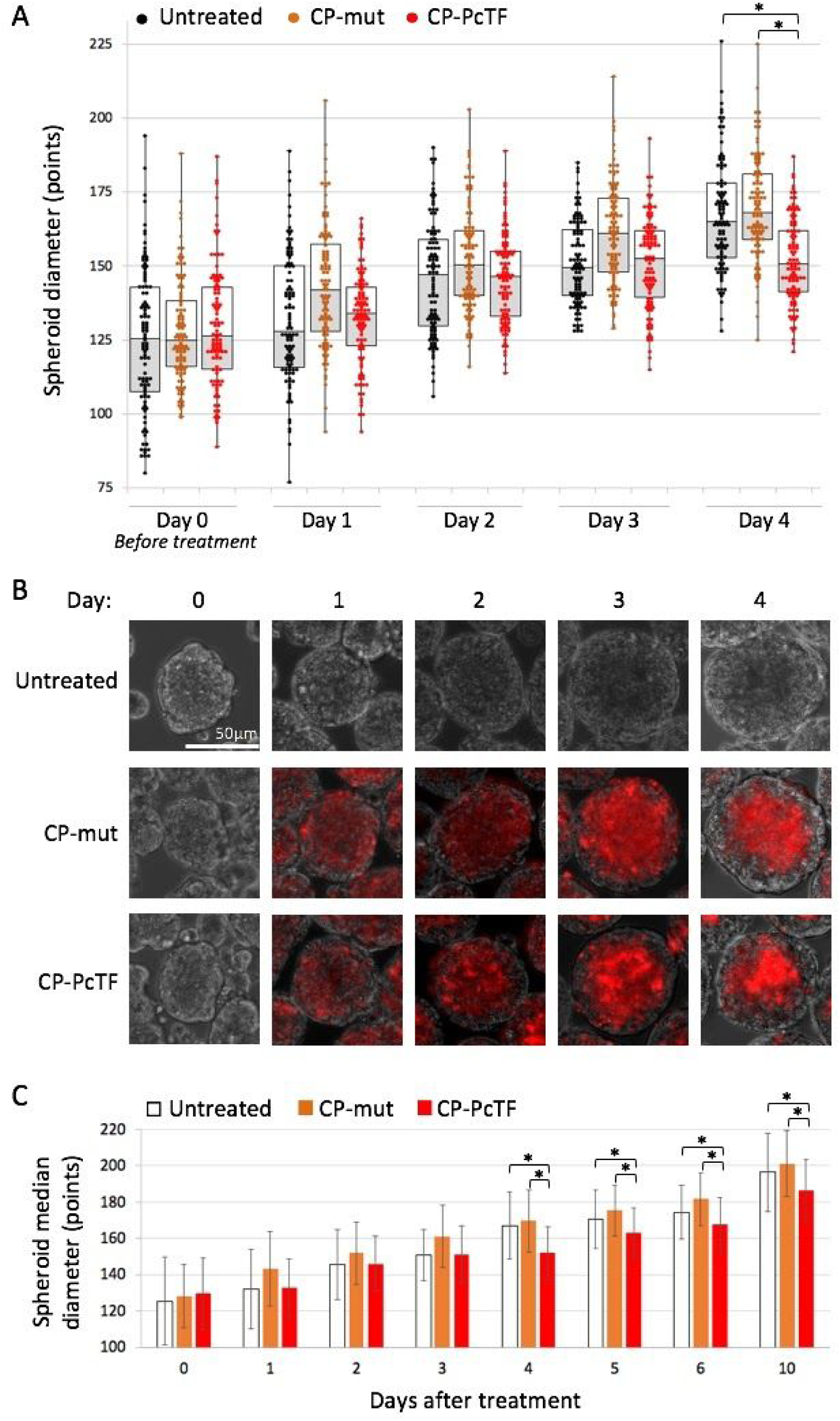
Comparisons of spheroid growth over time in CP-PcTF-treated samples versus controls. (A) The boxplot shows spheroid diameter values (center line, median; lower and upper boxes, 25th and 75th percentiles; lower and upper whiskers, minimum and maximum) for *n* = 100 spheroids per condition. Data were collected prior to treatment (day 0) and each day after treatment (days 1, 2, 3, and 4). (B) Representative images of U-2 OS spheroids used to generate the data shown in panel A. (C) The bar chart includes median values and standard deviations (thin bars) for the data from panel A (days 0 - 4) and additional time points (days 5, 6, and 10). Asterisks (*) denote *p* values < 0.001.

### RNA-seq analysis of CP-PcTF-treated cells showed modest effects on gene regulation

The imaging and flow cytometry results suggested highly efficient (> 86%) uptake of CP-PcTF in 2-D cultures. However, the RFP signal from the fusion protein was granular and not entirely within the nucleus. In our previous work, transgenically-expressed PcTF showed a homogeneous distribution of RFP signal entirely within the nucleus [18,28] (Figure 5A). These differences in subcellular distribution might affect the gene-regulating activity of the PcTF protein. Furthermore, PAGE analysis of purified CP-PcTF showed that at least half of the total protein may have been cleaved by endogenous protease activity in *E. coli*, [25]. Therefore, a substantial portion of CP-PcTF may have been truncated prior to delivery. To determine if the amount of CP-PcTF that may have entered the nucleus was sufficient to activate epigenetically repressed genes in the same manner as transgene-expressed PcTF, we treated 2-D U-2 OS cells with CP-PcTF or CP-mut (negative control protein) for 24 hours and harvested the RNA for next generation sequencing (RNA-seq).

**Figure 5.**
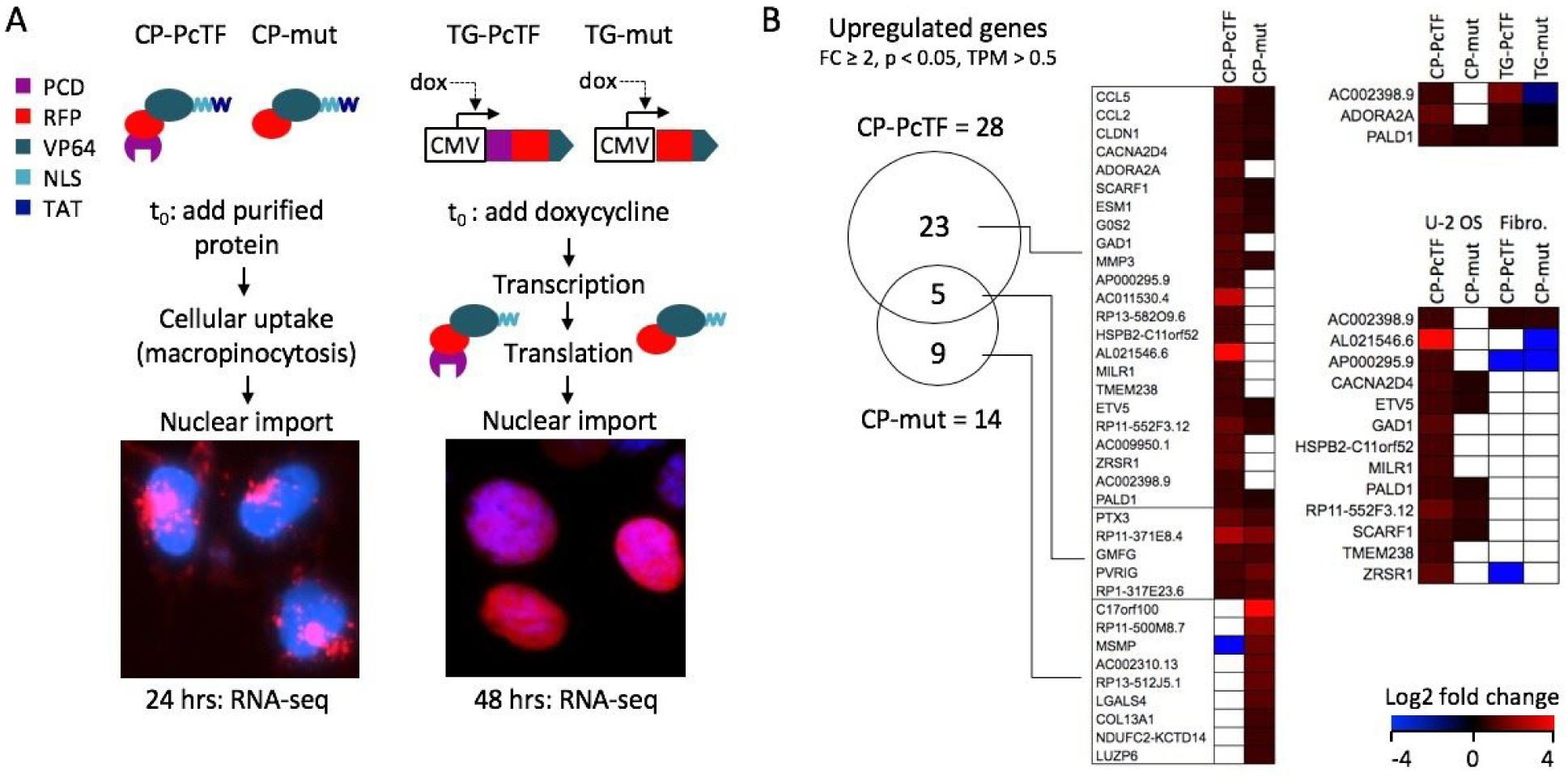
Gene expression changes in CP-PcTF-treated cells compared to transgenic PcTF-expressing cells. (A) Overview of TAT-mediated delivery of PcTF and the truncated negative control (mut) compared to dox-induced expression of the same fusion proteins (without TAT) from a stably integrated transgene (TG-PcTF or TG-mut). Images show RFP signals from the fusion proteins in U-2 OS cells stained with Hoescht (blue) DNA dye. (B) The Venn diagram shows shared and distinct genes with significant upregulation (fold change ≥ 2, *p* < 0.05, TPM > 0.5) for cells treated with CP-PcTF or the truncated control CP-mut. The heat maps compare log2 fold change values of genes in U-2 OS cells treated with cell-penetrating proteins (left), transgenic U-2 OS cells that express the fusion proteins (upper right), and U-2 OS versus non-cancer fibroblasts treated with cell-penetrating proteins (lower right). White boxes indicate *p* > 0.05 and/or TPM < 0.5.

Our RNA-seq analysis identified 28 genes that were significantly upregulated (fold change ≥ 2.0, *p* < 0.05) in CP-PcTF-treated versus untreated cells, and were expressed above the baseline value of 0.5 transcripts per kilobase million for both treated cell replicates (TPM > 0.5, see Methods for determination of threshold). Of these genes, 23 (92%) were upregulated by CP-PcTF and not by the negative control protein CP-mut (Figure 5B). Nine genes were exclusively upregulated in CP-mut, which lacks a targeting domain and is not expected to interact with genes. Three of the CP-PcTF-responsive genes, including *AC002398.9* (a locus that contains *PSENEN* and *LIN37*), *ADORA2A*, and *PALD1* were upregulated 3.7, 1.6, and 1.7-fold respectively by transgenic PcTF (TG-PcTF) and not by the TG-mut negative control in our previous RNA-seq study [18].

The upregulation of genes involved in the immune response (*CCL5, CCL2, MILR1*), and tumor suppression (*CLDN1, G0S2*) suggests that CP-PcTF might activate anti-proliferative and immunomodulatory gene expression in cancer cells. CP-PcTF might also affect gene expression in non-cancer cells because the cell penetrating peptide TAT on CP-PcTF is not cell-type specific, and the H3K27me3 mark (the binding target for PcTF) is virtually ubiquitous across cell types. To investigate off-target effects on non-cancer cells, we measured changes in gene expression in CP-protein-treated fibroblasts. Treatments with CP-PcTF and CP-mut were performed as described for U-2 OS. In CP-PcTF-treated fibroblasts 431 genes became significantly upregulated at least two-fold, and 187 (43%) of these were specifically activated CP-PcTF and not the control protein CP-mut. Although many more genes responded specifically to CP-PcTF in fibroblasts (187 in fibroblasts versus 23 in U-2 OS), 13 genes were uniquely upregulated in U-2 OS (Figure 5B). Overall, these results suggest that the CP-proteins used in this study might elicit a general, non-selective response, which underscores the importance of developing CP-proteins for cell-type-specific delivery or intracellular activity.

## DISCUSSION

This work represents the first characterization of a synthetic, cell penetrating chromatin sensor-actuator protein. Our previous work demonstrated that transgenic PcTF, which binds H3K27me3, activates epigenetically repressed genes in cancer cell lines including U-2 OS. In our current study, we demonstrated that the addition of a TAT signal supports efficient DNA-free delivery of purified PcTF protein to cells grown as monolayers and spheroids. We observed that transient delivery of cell-penetrating PcTF (CP-PcTF) inhibits the growth of U-2 OS spheroids to a modest but significant degree. CP-PcTF also has a modest effect on gene regulation compared to transgenically expressed PcTF.

We demonstrated that inclusion of the TAT signal within PcTF (CP-PcTF) facilitates very efficient uptake by U-2 OS cells in a concentration-dependent manner. Compared to 2-D monolayers, delivery of 0.1 to 1.0 μM was less efficient in 3-D spheroids. In spheroids treated with the highest concentration of CP-PcTF (1.0 μM), RFP signal was detected in only 45.4% of the cells. Our observation that uptake increased linearly with CP-PcTF dose suggests that delivery into spheroids could be improved, perhaps with prolonged or continuous treatment i.e., no washout of excess protein. Therefore, in future experiments it will be important to measure CP-PcTF uptake by individual cells in spheroids over time.

We showed that transient treatment with CP-PcTF temporarily inhibited the growth of U-2 OS spheroids. Spheroid size in CP-PcTF-treated samples was reduced four days after treatment compared to untreated cells and cells treated with a truncated version of CP-PcTF. From five to ten days, spheroid size continued to increase, but remained lower than the control values. It is possible the CP-PcTF affected only a subpopulation of cells within each spheroid since only 45.4% of the cells in spheroids treated with 1.0 μM CP-PcTF for 24 hours showed CP-PcTF uptake (RFP signal) in flow cytometry assays. Incomplete penetration CP-PcTF into each spheroid may have allowed unaffected cells to proliferate. Visualization of CP-PcTF uptake and markers for cell viability (e.g. staining for apoptosis and necrosis) in single cells within spheroids will help to determine the impact of CP-PcTF on cell proliferation.

Gene expression analysis revealed a modest level of gene activation in 2-D cultured cells that were treated with CP-PcTF. Based on our previous results from overexpression of PcTF from a transgene in U-2 OS cells [18,28], we expected CP-PcTF to enter the cell nucleus, bind multiple H3K27me3-enriched sites, and co-activate many epigenetically-repressed genes. We identified 103 genes that were upregulated specifically by CP-PcTF and not by the nonbinding control protein (CP-mut). Three of these genes were also specifically activated by transgene-expressed PcTF (TG-PcTF) in our previous study [18], which is just a small fraction of the 3807 genes that responded TG-PcTF. Although we observed nearly 100% uptake of CP-PcTF via flow cytometry at this concentration, high-resolution fluorescence microscopy suggested that inefficient nuclear import might hinder gene-regulating activity of the protein. Aggregates of RFP signal appeared near, and perhaps outside the nuclei of CP-PcTF-treated cells. Modifications to the fusion protein such as the addition of an endosomal escape signal and additional NLS tags (as recently demonstrated for Cas-based gene regulators proteins [7]) might improve nucleosomal import. Even if nuclear import were optimized, differences between transgene expression and protein delivery could influence PcTF’s gene-regulating activity. Continuous expression from TG-PcTF might produce steady state levels of PcTF, and therefore sustain transcriptional activation of H3K27me3-enriched target genes over time. In contrast, the delivery of a large concentration of CP-PcTF could result in a single, strong burst of transcription that becomes diminished as the CP-PcTF proteins are degraded or depleted during cell division. Transcription profiling by RNA-seq or RT-qPCR at additional time points after treatment would provide a clearer understanding of the dynamics of TG-PcTF and CP-PcTF-mediated gene regulation. Our gene profiling experiments also indicate that cell-specific targeting may be needed to prevent off-target effects of CP-PcTF. We observed that hundreds of genes became upregulated in non-cancer fibroblasts that were treated with CP-PcTF. The use of selective cell surface receptor-binding nanoparticles or protein ligands could help to avoid off-target toxicity.

In conclusion, this early demonstration of chromatin sensor-actuator proteins delivery into cells shows that the TAT signal can be used as a fusion protein module to support quick and robust protein uptake by U-2 OS cells in 2-D and 3-D cultures. Additional work is needed to optimize the yield of full-length TAT-tagged fusion proteins, and to achieve efficient endosomal escape and nuclear import of these proteins. Achieving these benchmarks might enable more robust CP-PcTF-mediated gene regulation in cancer cells, and enable the eventual development of a cancer-selective CP-PcTF.

## METHODS

### Construction of Plasmids for Recombinant Protein Expression

A recombinant plasmid insert encoding CP-PcTF was generated by consecutive PCRs using KAH87_MV10 as a template (https://benchling.com/hayneslab/f_/rmSYkAAU-synthetic-chromatinactuators-2-0). First, an amplicon that included the PcTF open reading frame (ORF) and a C-terminal nuclear localization signal (NLS), added a BamHI site (5’-GGATCC) to the 5’ end, and added a partial TAT signal (shown in brackets) to the 3’ end was generated via PCR using the following primers. The template-binding regions are underscored:

PcTF.BamHI.For: 5’-ATCA**GGATCC**<u>ATGGAGCTTTCAGCGG</u>

TAT.Rev.1: 5’-[CGTTTTTTACGACC]ATA<u>TACCTTGCGCTTTTTC</u>

The amplicon from this reaction was used as a template in a subsequent PCR reaction where the complete C-terminal TAT signal and a HindIII site (5’-AAGCTT) was added to the 3’ end of the PcTF-NLS ORF with the following primers. The TAT sequence is shown in brackets:

PcTF.BamHI.For: 5’-<u>ATCA</u>**<u>GGATCC</u>**<u>ATGGAGCTTTCAGCGG</u>

TAT.Rev.2: 5’-ATGCT**AAGCTT**CTA[ACGACGACGCTGACGA<u>CGTTTTTTACGACC]A</u>

The PCR product was digested with BamHI and HindIII, purified on a PCR cleanup column (Qiagen 28106), and ligated into BamHI/ HindIII-linearized and gel-purified pET28. CP-mut was cloned by PCR amplification of CP-PcTF with the following primers:

McVP64.BamHI.For: 5’-GGTCGC**GGATCC**<u>GTGAGCAAGGGCGAGGAG</u>

TAT.Rev.2: 5’-<u>ATGCT</u>**<u>AAGCTT</u>**<u>CTA[ACGACGACGCTGACGACGTTTTTTACGACC]A</u>

The CP-mut amplicon was digested and ligated into pET28 as described for CP-PcTF.

### Expression and Purification of Recombinant Proteins

Proteins were expressed, purified, and quantified as described in Tekel et. al. 2018 [20].

### Cell Culture

U-2 OS cells (ATCC HTB-96) were maintained in McCoy’s 5A (Modified) Medium supplemented with 10% (v/v) fetal bovine serum (FBS) and 1% (v/v) penicillin-streptomycin (all from ThermoFisher). Cells were maintained in a 37°C incubator with 5% CO_2_ and passaged once ∼80% confluent. 2-D cultures were seeded with 250,000 cells per well in a 6-well tissue culture-treated plate and grown for 24 hours prior to treatment (About 30% confluency). 3-D spheroids were generated by seeding 2.0E6 cells per well of a 6-well cell repellent plate (VWR) in a final volume of 4.0 mL complete media and placed on an orbital shaker at 95 RPM for 24 hours. Half media change was performed every 24 hours. Spheroids were cultured for a minimum of 48 hours before treatment. BJ fibroblasts (ATCC CRL-2522) were maintained in Dulbecco’s modified eagle medium supplemented with 10% (v/v) fetal bovine serum (FBS) and 1% (v/v) penicillin-streptomycin (ThermoFisher). Cells were maintained in a 37°C incubator with 5% CO_2_ and passaged once ∼80% confluent.

### Microscopy and Flow Cytometry

Fluorescent images of each well at 10x and 20x were obtained using an EVOS FL Cell Imaging System (Invitrogen, model AMF4300) using the RFP channel (excitation/emission: 470/510 nm) with a 500 ms exposure time. Prior to flow cytometry, cells from 2-D or 3-D cultures were harvested from one well of a 6-well plate and dissociated with Accutase (Thermo Fisher #A1110501), washed three times with 1x PBS, resuspended in 1x PBS, and analyzed using a BD Accuri C6 cytometer (BD Biosciences). RFP-positive cells were measured using the “FL-3” channel (excitation/emission: 488/670 nm) and gated using untreated cells as background. Data was analyzed with FlowJo software (v10).

### 3-D Spheroid Growth Analysis

Five independent 10x phase contrast images were taken for each condition on each day using an EVOS FL inverted fluorescence microscope (Invitrogen, model AMF4300). Cell diameter (in points) was measured in Adobe Illustrator CS6 for 100 spheroids per condition per day. *P* values were calculated using a Student’s two-tailed, homoscedastic t-test.

### RNA-seq Assay and Data Analysis

Total RNA was extracted from duplicate samples (1.5 million cells per sample) for each experimental condition. RNA extraction was performed using Nucleospin RNA (Macherey-Nagel 740955.250). RNA-seq was performed by the next generation sequencing core at Arizona State University (ASU). Total RNA was ribo-depleted (KAPA Ribo-Zero RNA HyperPrep Kit with RiboErase HMR, Roche #KK8560), sheared to roughly 250 bp, and then converted to cDNA. Illumina-compatible adapters with unique indexes (IDT #00989130v2) was then ligated onto each sample individually. The adapter ligated molecules were then cleaned using AMPure beads (Agencourt Bioscience/Beckman Coulter, A63883), and amplified with Kapa’s HIFI enzyme (KK2502). Each library was analyzed for fragment size on an Agilent Tapestation, and quantified by qPCR (KAPA KK4835) on a Quantstudio 5 instrument (Thermo Fisher) before multiplex pooling and single-end sequencing on a 1×75 flow cell on the NextSeq500 platform (Illumina). Total reads were 33 to 65 million per sample.

Differential expression and normalization were performed with edgeR software [29]. Transcripts per kilobase million (TPM) for each gene was calculated as reads per kilobase gene length (RPK) divided by a reads per million scaling factor (the sum of all RPK’s divided by 1.0E6) [30]. We selected 0.5 as the minimum TPM for gene expression, based on the default threshold for the EMBL-EBI Expression Atlas (https://www.ebi.ac.uk/gxa/FAQ.html). TPM’s of genes known to be silenced in U-2 OS [31] were consistently near or less than 0.5 in our untreated U-2 OS sample duplicates: *CDKN2A* (0.04, 0.08), *CDKN2B* (0.50, 0.53), *HIC1* (0.13, 0.51), *DLX5* (0.18, 0.13). The TPM’s of three active housekeeping genes [18] were above 0.5: *GAPDH* (3951.50, 4692.36), *CHMP2A* (18.86, 20.16), *ACTB* (2938.25, 3612.29). The Venn diagram was generated from gene symbol lists and the Venny 2.1 tool online (http://bioinfogp.cnb.csic.es/tools/venny/) [32].

## AUTHOR CONTRIBUTIONS

SJT and KAH conceptualized the project. SJT cloned, expressed, purified, quantified recombinant proteins, and prepared total RNA from treated U-2 OS cells. NB performed cell culture. SJT and NB performed microscopy and flow cytometry. KAH supervised the experiments and performed gene expression comparisons. SJT and KAH generated figures, analyzed data, and wrote the manuscript.

## ACKNOWLEDGEMENTS

We thank Jason Steele and the Arizona State University Bioinformatics Core for their help with RNA quality analysis, RNA sequencing, and RNA-seq data processing. Peptides were a generous gift from Tara MacCulloch in the Nick Stephanopolous lab. We also thank David Brafman for reagents and supplies.

## REFERENCES

1. Du S, Liew SS, Li L, Yao SQ. Bypassing Endocytosis: Direct Cytosolic Delivery of Proteins. J Am Chem Soc. 2018;140: 15986–15996.

2. Leader B, Baca QJ, Golan DE. Protein therapeutics: a summary and pharmacological classification. Nat Rev Drug Discov. 2008;7: 21–39.

3. Rees HA, Komor AC, Yeh W-H, Caetano-Lopes J, Warman M, Edge ASB, et al. Improving the DNA specificity and applicability of base editing through protein engineering and protein delivery. Nat Commun. 2017;8: 15790.

4. Kintzing JR, Filsinger Interrante MV, Cochran JR. Emerging Strategies for Developing Next-Generation Protein Therapeutics for Cancer Treatment. Trends Pharmacol Sci. 2016;37: 993–1008.

5. Zelikin AN, Ehrhardt C, Healy AM. Materials and methods for delivery of biological drugs. Nature Chemistry. 2016. pp. 997–1007. doi: 10.1038/nchem.2629

6. Zuris JA, Thompson DB, Shu Y, Guilinger JP, Bessen JL, Hu JH, et al. Cationic lipid-mediated delivery of proteins enables efficient protein-based genome editing in vitro and in vivo. Nat Biotechnol. 2015;33: 73–80.

7. Staahl BT, Benekareddy M, Coulon-Bainier C, Banfal AA, Floor SN, Sabo JK, et al. Efficient genome editing in the mouse brain by local delivery of engineered Cas9 ribonucleoprotein complexes. Nat Biotechnol. 2017;35: 431–434.

8. Woo JW, Kim J, Kwon SI, Corvalán C, Cho SW, Kim H, et al. DNA-free genome editing in plants with preassembled CRISPR-Cas9 ribonucleoproteins. Nat Biotechnol. 2015;33: 1162–1164.

9. Okamoto Y, Kojima R, Schwizer F, Bartolami E, Heinisch T, Matile S, et al. A cell-penetrating artificial metalloenzyme regulates a gene switch in a designer mammalian cell. Nature Communications. 2018. doi: 10.1038/s41467-018-04440-0

10. Krosl J, Austin P, Beslu N, Kroon E, Humphries RK, Sauvageau G. In vitro expansion of hematopoietic stem cells by recombinant TAT-HOXB4 protein. Nat Med. 2003;9: 1428–1432.

11. Michiue H, Tomizawa K, Wei F-Y, Matsushita M, Lu Y-F, Ichikawa T, et al. The NH2 terminus of influenza virus hemagglutinin-2 subunit peptides enhances the antitumor potency of polyarginine-mediated p53 protein transduction. J Biol Chem. 2005;280: 8285–8289.

12. Guidotti G, Brambilla L, Rossi D. Cell-Penetrating Peptides: From Basic Research to Clinics. Trends Pharmacol Sci. 2017;38: 406–424.

13. Essafi M, Baudot AD, Mouska X, Cassuto J-P, Ticchioni M, Deckert M. Cell-penetrating TAT-FOXO3 fusion proteins induce apoptotic cell death in leukemic cells. Mol Cancer Ther. 2011;10: 37–46.

14. Kwon YD, Oh SK, Kim HS, Ku S-Y, Kim SH, Choi YM, et al. Cellular manipulation of human embryonic stem cells by TAT-PDX1 protein transduction. Mol Ther. 2005;12: 28–32.

15. Kelly TK, De Carvalho DD, Jones PA. Epigenetic modifications as therapeutic targets. Nature Biotechnology. 2010. pp. 1069–1078. doi: 10.1038/nbt.1678

16. Baskin NL, Haynes KA. Chromatin engineering offers an opportunity to advance epigenetic cancer therapy. Nat Struct Mol Biol. 2019;26: 842–845.

17. Tekel SJ, Haynes KA. Molecular structures guide the engineering of chromatin. Nucleic Acids Res. 2017;45: 7555–7570.

18. Nyer DB, Daer RM, Vargas D, Hom C, Haynes KA. Regulation of cancer epigenomes with a histone-binding synthetic transcription factor. NPJ Genom Med. 2017;2. doi: 10.1038/s41525-016-0002-3

19. Olney KC, Nyer DB, Vargas DA, Wilson Sayres MA, Haynes KA. The synthetic histone-binding regulator protein PcTF activates interferon genes in breast cancer cells. BMC Syst Biol. 2018;12: 83.

20. Tekel SJ, Vargas DA, Song L, LaBaer J, Caplan MR, Haynes KA. Tandem Histone-Binding Domains Enhance the Activity of a Synthetic Chromatin Effector. ACS Synth Biol. 2018;7: 842–852.

21. Wadia JS, Stan RV, Dowdy SF. Transducible TAT-HA fusogenic peptide enhances escape of TAT-fusion proteins after lipid raft macropinocytosis. Nat Med. 2004;10: 310–315.

22. Lee Y-J, Erazo-Oliveras A, Pellois J-P. Delivery of macromolecules into live cells by simple co-incubation with a peptide. Chembiochem. 2010;11: 325–330.

23. Akishiba M, Takeuchi T, Kawaguchi Y, Sakamoto K, Yu H-H, Nakase I, et al. Cytosolic antibody delivery by lipid-sensitive endosomolytic peptide. Nat Chem. 2017;9: 751–761.

24. Lönn P, Kacsinta AD, Cui X-S, Hamil AS, Kaulich M, Gogoi K, et al. Enhancing Endosomal Escape for Intracellular Delivery of Macromolecular Biologic Therapeutics. Sci Rep. 2016;6: 32301.

25. Patel SG, Sayers EJ, He L, Narayan R, Williams TL, Mills EM, et al. Cell-penetrating peptide sequence and modification dependent uptake and subcellular distribution of green florescent protein in different cell lines. Sci Rep. 2019;9: 6298.

26. Yang Y, Ma J, Song Z, Wu M. HIV-1 TAT-mediated protein transduction and subcellular localization using novel expression vectors. FEBS Lett. 2002;532: 36–44.

27. De Luca A, Raimondi L, Salamanna F, Carina V, Costa V, Bellavia D, et al. Relevance of 3d culture systems to study osteosarcoma environment. J Exp Clin Cancer Res. 2018;37: 2.

28. Haynes KA, Silver PA. Synthetic Reversal of Epigenetic Silencing. Journal of Biological Chemistry. 2011. pp. 27176–27182. doi: 10.1074/jbc.c111.229567

29. Robinson MD, McCarthy DJ, Smyth GK. edgeR: a Bioconductor package for differential expression analysis of digital gene expression data. Bioinformatics. 2010;26: 139–140.

30. Li B, Dewey CN. RSEM: accurate transcript quantification from RNA-Seq data with or without a reference genome. BMC Bioinformatics. 2011. doi: 10.1186/1471-2105-12-323

31. Easwaran H, Johnstone SE, Van Neste L, Ohm J, Mosbruger T, Wang Q, et al. A DNA hypermethylation module for the stem/progenitor cell signature of cancer. Genome Res. 2012;22: 837–849.

32. Collazos JCO. Venny 2.1.0. [cited 15 Jan 2020]. Available: https://bioinfogp.cnb.csic.es/tools/venny/index.html

