## Supplemental for "Delivery of cell-penetrating chromatin sensor-actuators to human osteosarcoma cells"

Table S1. Peptides that were incubated with PcTF prior to delivery to U-2 OS cells.

Figure S1. Images of cells treated for 24 hours with PcTF plus each cell penetrating peptide.

Figure S2. PAGE analysis of column-purified CP-PcTF proteins.

Supplemental Methods

Supplemental References

|  | **Name** | **Primary structure** | **Reference** |
| --- | --- | --- | --- |
| Peptide 1 | TAT-HA2 | RRRQRRKKRGGDIMGEWGNEIFGAIAGFLG | Wadia 2004 |
| Peptide 2 | E5-TAT | GLFEAIAEFIENGWEGLIEGWYGGRKKRRQRRR | Lee 2010 |
| Peptide 3 | TAT | GRKKRRQRRR | Lee 2010,  Wadia 2004 |
| Peptide 4 | L17E | WLTALKFLGKHAAKHEAKQQLSKL-amide | Akishiba 2017 |

**Table S1.** Peptides that were incubated with PcTF prior to delivery to U-2 OS cells.

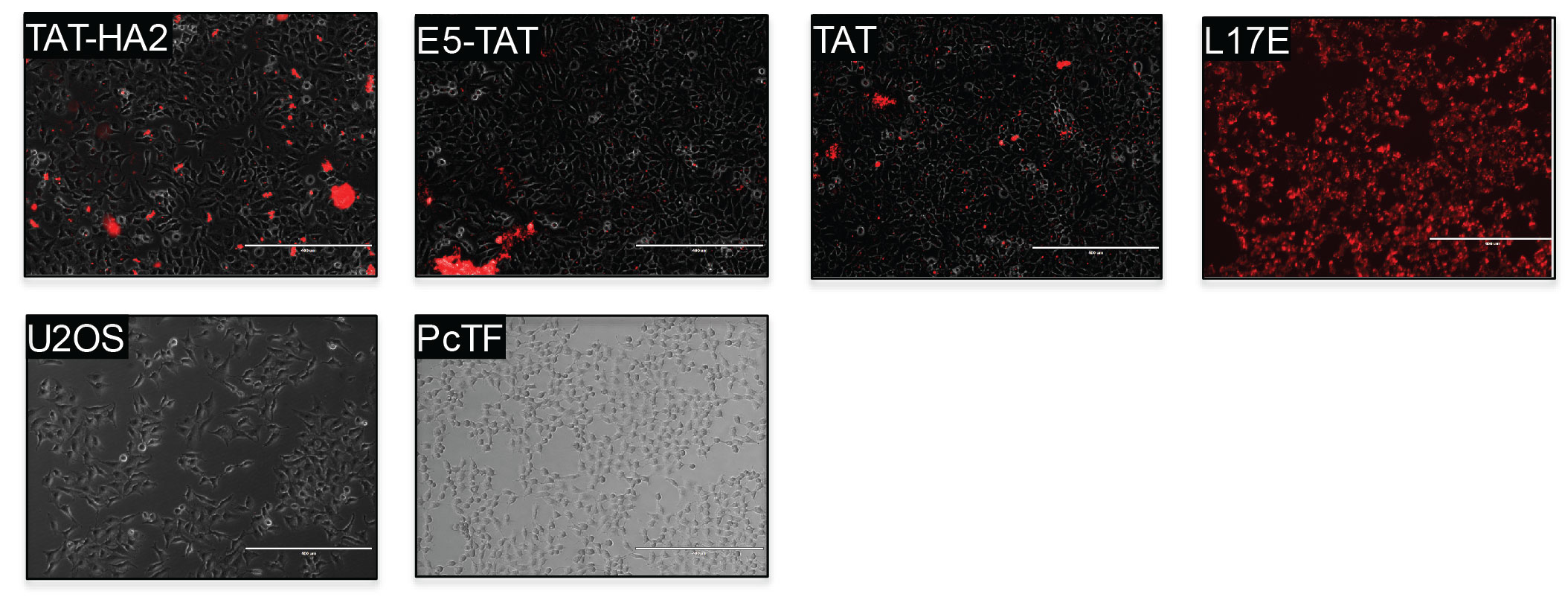

**Figure S1.** Images of cells treated for 24 hours with PcTF plus each cell penetrating peptide. Cells were treated with 1.0 μM PcTF plus 5.0 μM of each cell penetrating peptide for 24 hours, washed once with 1x PBS and imaged for mCherry red fluorescent protein signal (RFP). U2OS = untreated cells, PcTF = cells treated with PcTF only.

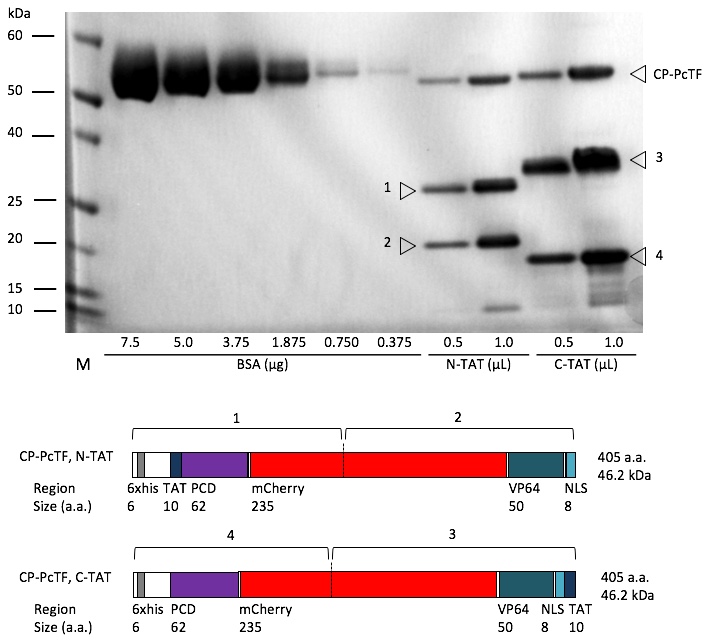

**Figure S2.** PAGE analysis of column-purified CP-PcTF proteins. Fusions that carried the TAT signal at either the N-terminus (N-TAT) or the C-terminus (C-TAT) were expressed and purified as described in Methods. Samples for denaturing polyacrylamide gel electrophoresis (PAGE) were prepared by heating purified, concentrated protein and ddH_2_O (to 9 μL total) plus 3 μL 4x NuPAGE LDS Sample Buffer (Thermo Fisher #NP0007) at 100°C for 5 min. Bovine Serum Albumin (BSA) samples were prepared the same way by diluting a stock of BSA in water. Samples (cooled to room temperature) and a pre-stained protein marker “M” (10 μL, Thermo Fisher #10748010) were electrophoresed at 200 V in a 4-12% Bis-Tris gel (Thermo Fisher #NP0322BOX) with 1x NuPAGE MOPS-SDS buffer (Thermo Fisher NP001) in an XCell SureLock vertical chamber (Invitrogen #EI0001). The gel was stained with coomassie dye R-250 (Imperial Protein Stain, Thermo Fisher #24615) overnight, destained in dH_2_O for 6 hrs., stained again for 2 hrs., destained overnight, and analyzed on a PXi4 imager (Syngene). A standard curve, based on signals from the BSA lanes, was calculated using GeneSys software (Syngene). Maps below the PAGE image show the approximate hypothetical site of cleavage (vertical dotted line) within mCherry that may have produced the two smaller bands in each lane for CP-PcTF N-TAT (bands 1 and 2) and C-TAT (bands 3 and 4). The shifts in band sizes, due to the different positions of the TAT signal, are consistent with this hypothesis. Fragments 1 and 3 carry the 6x his tag and are therefore expected to co-purify with full length CP-PcTF. Fragments 2 and 4 may have co-purified due to the addition of a second C-terminal 6xhis tag (in-frame with the cloned ORF in pET28) via read-through past the single stop codon.

**SUPPLEMENTAL METHODS**

**Peptide Synthesis**

Peptides 1-4 (Table S1) were synthesized on a resin using standard Fmoc chemistries. Analysis of the peptides by analytical HPLC and MALDI-TOF confirmed that the peptides had the correct expected masses. Peptides were brought to a final concentration of 500 μM in dH_2_O.

**Preparation of of PcTF-peptide Complexes and Delivery to Cells**

U-2 OS cells were grown to 30% confluency in 12-well tissue culture-treated plates under 1.0 mL complete growth medium. PcTF was premixed with Peptide 1, 2, 3, or 4 (Table S1) in a final volume of 500 μL complete growth medium, incubated at room temperature for 10 min., and added directly to U-2 OS cells at the following final concentrations in 1.0 mL growth medium: 5.0 μM peptide plus 1.0 μM PcTF, 1.0 μM PcTF only, or 5.0 μM peptide only. Cells were incubated for 24 hours before removing media and washing with 1.0 mL 1x PBS three times, and imaged via fluorescence microscopy (Invitrogen EVOS AMF4300).

**SUPPLEMENTAL REFERENCES**

1. Wadia JS, Stan RV, Dowdy SF. Transducible TAT-HA fusogenic peptide enhances escape of TAT-fusion proteins after lipid raft macropinocytosis. Nat Med. 2004; 10: 310-315.
2. Lee Y-J, Erazo-Oliveras A, Pellois J-P. Delivery of macromolecules into live cells by simple co-incubation with a peptide. Chembiochem. 2010; 11(3): 325-330.
3. Akishiba M, Takeuchi T, Kawaguchi Y, Sakamoto K, Yu HH, Nakase I, Takatani-Nakase T, Madani F, Gräslund A, Futaki S. Cytosolic antibody delivery by lipid-sensitive endosomolytic peptide. Nat Chem. 2017; 9(8):751-761.
